# Western Ghats Myrtaceae are not Gondwana elements but likely dispersed from south-east Asia

**DOI:** 10.1101/2020.04.12.037960

**Authors:** Rajasri Ray, Balaji Chattopadhyay, Kritika M Garg, TV Ramachandra, Avik Ray

## Abstract

Gondwana break-up is one of the key sculptors of the global biogeographic pattern including Indian subcontinent. Myrtaceae, a Gondwana family, demonstrates a high diversity across Indian Western Ghats. A rich paleo-records in Deccan Intertrappean bed in India strive to reconcile two contending hypothesis of origin and diversification of Myrtaceae in India, namely Gondwana elements floated on peninsular India or lineages invading from south-east Asia. In this study, we have reconstructed the biogeographic history of Myrtaceae of Western Ghats of India combining fossil-calibrated phylogeny, ancestral area reconstruction, and comparing various dispersal models.

Phylogenetic reconstructions divided lineages into multiple clades at the tribal level that mostly is in agreement with previous descriptions. Dated tree indicated a presumable Gondwanan origin of Myrtaceae (95.7 Ma). The Western Ghats taxa are relatively younger and seem to have arrived either through long-distance oceanic dispersal in middle Eocene (*Rhodomyrtus tomentosa*), and Miocene (*Eugenia*) or through major overland dispersals throughout Miocene (*Syzygium*). However, late Cretaceous fossilsin India lends support to the existence of proto-Myrtaceae on floating peninsular India. Taken together, we conclude that Myrtaceae lineages on the biotic ferry were perhaps extirpated by Deccan volcanism leaving a little or no trace of living Gondwanan link, while the Western Ghats was likely enriched by dispersed elements either from south-east Asia or from South America.

---

India has a unique yet complex geologic origin as aptly reflected in its rich biological legacy. A fraction of diversity is attributed to Gondwana elements whereas the rest consisting of dispersed biota with Sino-Tibetan, Indo-Malayan, or Mediterranean affinities (Mani, 1974). The Western Ghats, a continuous mountain chain spanning 1500 km^2^ along the western coast of India, spread across five states from 8°N to 22°N and 73°E to 77°E. Being a global hotspot, it exhibits rich floral diversity with more than 8000 taxa of flowering plants, including 7402 species, 117 subspecies, and 476 varieties (Nayar 2014, CEPF 2007). Distinct temperature and rainfall gradients prevail across south-north and west-east directions of the Ghats. It influences the distribution of biota that is discernible in species richness and endemicity patterns (Gimaret-Carpentier et al. 2003). The southern part, presumed to be refugia, is particularly rich in Gondwana flora and could be a home of ancient lineages (Prasad et al. 2009).

Myrtaceae, the eighth largest flowering plant family of more than 145 genera and 5500 species (Govaerts et al. 2009), is of Gondwanan origin (Thornhill et al. 2015). It is represented by 17 tribes mostly in Southern Hemisphere, and quite widespread with centres of diversity in Australia, south and south-east Asia, tropical to southern temperate America, but has little representation in Africa (Sytsma et al. 2003). In India, taxa from tribes, Syzygieae and Myrteae, overwhelm Myrtaceae diversity. There are almost 26 species (1 *Meteromyrtus* sp., 9 *Eugenia* spp. and 16 *Syzygium* spp., 1 *Rhodomyrtus*) reported in Western Ghats (henceforth WG) region (Pascal & Ramesh, 1997); though taxonomic discrepancy constrains species delimitation (Byng et al., 2015; Sujanapal and Kunhikannan 2017). In addition, several species of *Syzygium* and *Eugenia* also abound the north-eastern region of India. Apart from a large number of endemic lineages, a few were introduced and have been cultivated for various reasons, such as, in afforestation, as fruit-crop or ornamental (e.g. *Eucalyptus, Callistemon, Melaleuca, and Psidium*) (Pascal and Ramesh 1997; Nayar et al. 2006).

The presence of disjunct Myrtaceae flora across WG and north-eastern India is intriguing. It evokes two alternate routes in terms of their origin and evolution: the first route posits that proto-Myrtaceae remained on the Indian raft after Gondwana vicariance (biotic ferry or Noah’s ark model) (Conti et al. 2002) and colonised WG later (figure 1a - c). Had they persisted, these lineages would be expected to emerge from deeper phylogenetic nodes. The abundance of fossils in Deccan Intertrappean Bed and elsewhere in India provides support to this Gondwana lineage hypothesis (Srivastava, 2011 and references therein; table S1). In contrast, the alternate hypothesis posits floral dispersals from south-east Asia (originally from the part of Gondwana that now makes up Australia) which eventually led Myrtaceae lineages to reach WG (Mani, 1974). This late arrival via south-east Asia ascribes it to relatively younger taxa (figure 1d).

**Figure 1:**
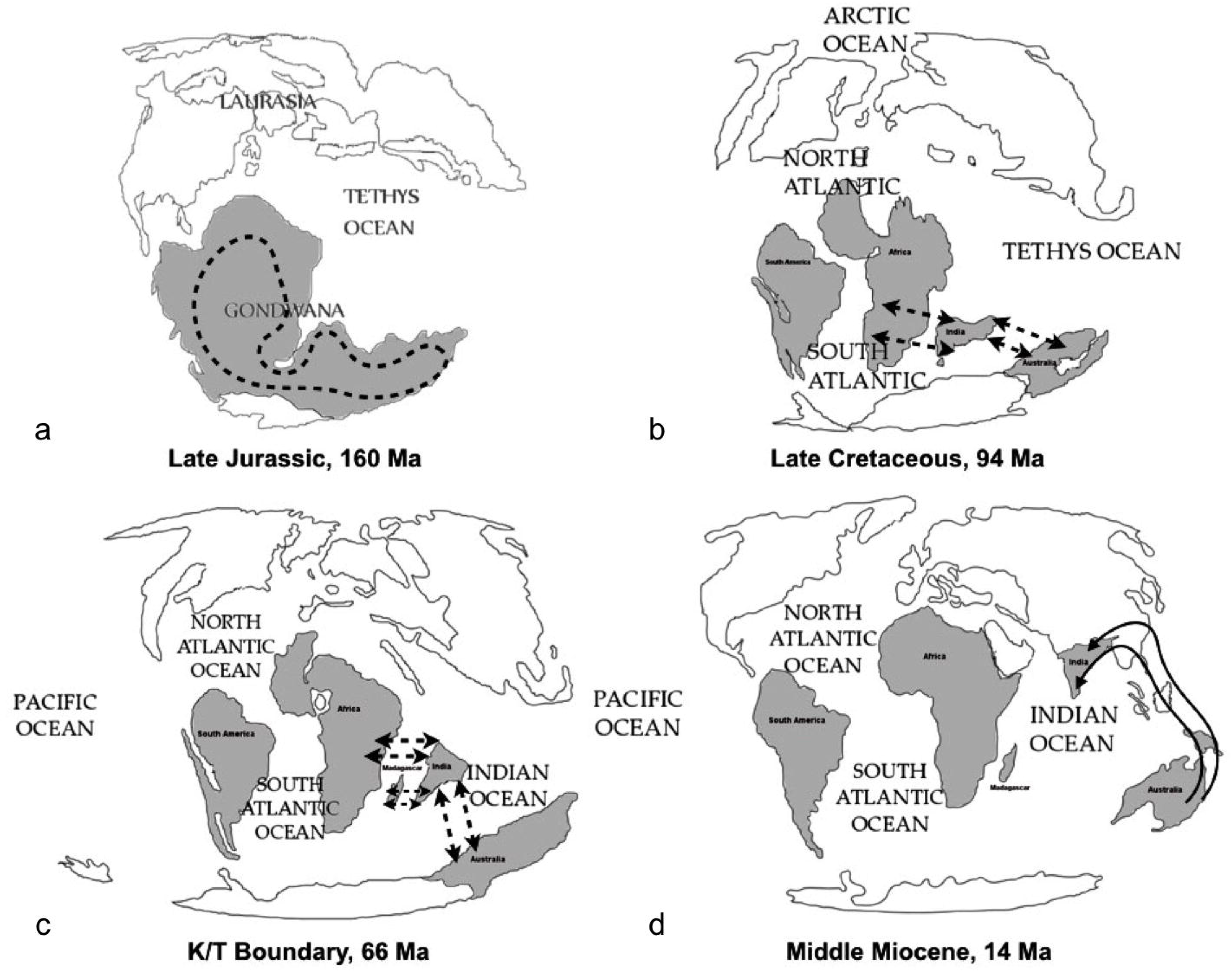
Evolutionary trajectory of Indian Myratceae from a. Late Jurassic: showing putative distribution of Proto-Myrtaceae, b. Late Cretaceous and c. K/T boundary: Movement of Indian peninsular mass and biotic exchange with Afro-Madagascar, Austrlia, d. Middle Miocene: India sutured to mainland Asia and overland dispersal from Australia and south-east Asia (redrawn after Scotese 2001)

However, none of these hypotheses have been rigorously tested within a phylogenetics driven biogeography framework and the origin of Indian Myrtaceae remains unsolved. We have sought to address this question by generating sequence data from a host of Indian Myrtaceae species and tested the following hypotheses: i) whether the extant lineages stem from Gondwana, or, ii) they represent dispersed elements, or, iii) both vicariance and dispersal have co-contributed to the generation of modern Myrtaceae diversity in WG. We performed fossil calibrated divergence dating, compared various dispersal-vicariance scenarios to reconstruct the history of Western Ghats Myrtaceae.

## 1. Materials and methods

### 1.1 Taxon sampling, DNA extraction, amplification, and sequencing

Relatively fewer taxonomic studies on Indian Myrtaceae impeded exhaustive taxon sampling. Moreover, inadequate information on species distribution (e.g. *S. agasthyamalayanum, S. beddomei, S. microphyllum* and many others) coupled with erroneous identification owing to a few characters were other obstacles (Sujanapaul and Kunhikannan 2017). Hence, in order to circumvent this problem, we sampled fifteen species of Myrtaceae from the WG of India(13 species of *Syzygium*, 1 species each of *Eugenia* and *Rhodomyrtus sp*) with little or no taxonomic disputes. Additionally, we supplemented our dataset with seventy additional species from GenBank; so that the final dataset consisted of a total of 81 species of Myrtaceae *sensu lato* (Wilson et al. 2005) as ingroup and four species (*Crypteronia paniculata, Alzatea verticillata, Olinia ventosa, Penaea mucronata*) from Crypteroniaceae, and Peneaceae as outgroup following Sytsma et al. (2003). Voucher specimens information and corresponding GenBank accession numbers of the existing and the new taxa are enlisted in table-S2-3.

We extracted DNA following modified Doyle and Doyle (1987), amplified the regions of chloroplast (*matk*), mitochondria (*ndhF*), and nucleus (ITS) using the standard protocol, and sequenced (for primer sequences, see refs. Biffin et al. 2006 and table S4). We aligned the sequences using ClustalX version 1.83 (65) and adjusted manually in MEGA 5.2 (Tamura et al. 2011).

### 1.2 Dated Molecular Phylogeny and fossil calibration

#### a) Molecular phylogeny

Sequences were concatenated using Sequence Matrix v1.7.8 (Vaidya et al., 2011) and data partition analyses were performed in Partition Finder v1.1.1 (Lanfear et al., 2012) using AIC method for model selection. We used unlinked branch lengths and greedy search scheme. Seven partitions for the data, consisting of a single partition for ITS and 3 partitions each for both *matk* and *ndhF* based on codon position, were defined; and finally, the best partition scheme determined by partition finder used to define data partitions for all concatenated phylogenetic reconstructions. We performed phylogenetic tree reconstructions with concatenated data using maximum likelihood method in RaxML GUI v1.5 (Silvestro and Michalak, 2012) and Bayesian reconstruction in MrBayes. We used the most complex model of sequence evolution (GTR◻+◻gamma) in RaxML. After performing a hundred runs implementing a thorough bootstrap algorithm with 1000 replicates, finally, we have obtained a maximum likelihood (ML) tree. In MrBayes, we performed two independent runs each for 2,000,000 generations with four chains keeping the swapping frequency and temperature at default values. We sampled the trees at every 500th generation and obtained diagnostics at every 5,000th generation. In course, we ensured that the standard deviation of the split frequency has reached below 0.01 by the end of each run. Finally, we checked the parameter convergence in Tracer v1.6 (Rambaut and Drummond 2007) and obtained the consensus trees from MrBayes after discarding 25 % of the trees as burn-in.

In addition, we also performed species tree reconstruction under Bayesian inference in *BEAST v1.8.2 (Drummond et al., 2012) considering each locus as a separate partition. For each locus, the substitution model was determined through jModelTest v 2.1.7 (Guindon and Gascuel, 2003; Darriba et al., 2012), and a separate relaxed molecular clock model for each locus was adopted to estimate relative clock rates. We implemented Yule speciation process with random starting gene trees and conducted two independent runs (of 100 million generations each and sampling every 100th generation) followed by a checking of parameter convergence of both the runs in Tracer (Rambaut et al., 2014). We combined the two runs were combined in Log Combiner 1.8.2 with a 50% burn-in while resampling at a frequency of 1000 generations. Subsequently, a maximum clade credibility tree was constructed using median heights in Tree Annotator 1.8.3 while allowing for a further 20% burn-in using a posterior probability limit of 0.95.

#### b) Fossil calibration and divergence dating

In addition to phylogenetic reconstruction, we also used the partitioned dataset of species tree reconstruction to obtain dates of divergences of various nodes in BEAST v1.8.3 (Drummond et al. 2012). We performed two independent runs of 50 million generations each, with sampling at every 1000th generation. We used the Birth-Death process of speciation with random starting trees and four time points for calibration of divergence dates based on previous studies of Biffin et al. (2010), Shukla et al. (2012) and Thornhill et al. (2015) (table - 1).

**Table-1:** Fossil calibrations used for molecular dating of major nodes of Myrtaceae. All priors are normally distributed.

| | Prior mean ± SD | Reference |
| --- | --- | --- |
| Myrtaceae stem | $92 \pm 3.5$ | Biffin et al. (2010) |
| Myrtaceae crown | $85 \pm 3.5$ | Thornhill et al. (2015) |
| Myrtoideae crown | $71.5 \pm 3$ | Thornhill et al. (2015) |
| Eucalypteae stem | $65 \pm 7$ | Shukla et al. (2012) |

Despite a continued effort to unravel historical biogeography of Myrtaceae, the results tended to vary greatly partly due to different fossil dates (Vasconcelos et al. 2017). To avoid this, we largely followed the fossil dates in the previous studies by Biffin et al. (2010) and Thornhill et al. (2015). Biffin et al. (2010) used the pollen taxon *Myrtaceidites lisamae* recorded from fossils obtained from multiple places, e.g. Gabon (Herngreen, 1975; Boltenhagen, 1976 Muller, 1981), Borneo (Muller, 1968), and from Colombia (van der Hammen, 1954) and that provided the earliest estimate of Myrtaceae radiation. Hence, we calibrated the Myrtaceae stem following this estimate at 92 Ma (± 3.5).

We calibrated Myrtaceae [85 (± 3.5) ma] and Myrtoideae [71.5 (± 3)] crown dates adhering to the large-scale phylogenetic study by Thornhill et al. (2015). They obtained 85 Ma (75.0–93.2 Ma) as the crown date of Myrtaceae when the two subfamilies diverged in the Late Cretaceous and Myrtoideae crown dates as 71.5 Ma (65.4-78.3) when tribes Xanthostemoneae, Osbornieae, and Lophostemoneae split from the rest of the Myrtoideae.

Lastly, a Eucalypt fossil, *Eucalyptus ghughuensis,* found in Deccan Intertrappean Bed (henceforth DIB) from India was used to calibrate the Eucalypteae stem in our phylogeny (Shukla et al. 2012).DIB of India has luxuriant records of Myrtaceae fossils preserved over a long period from Danian-Maastrichtian. However, the xylotomical features of the fossil, e.g. characteristic solitary vessels, paratracheal and apotracheal parenchyma, vasicentric tracheids, fine rays highlighted its morphological similarity with both Fagaceae and Myrtaceae. However, a characteristic echelon arrangement of the vessels has suggested a Myrtaceae affinity, ascribing to *Eucalyptus, Melaleuca,* or *Xanthostemon.* The final inclusion into the genus *Eucalyptus* was based on semi-ring porous wood which is absent in other two genera but present in a few species of *Eucalyptus*. It is the oldest known Myrtaceae fossil not only from India perhaps in the world.

For all calibrations, priors were provided with the normal distribution. Parameter convergence of each run was confirmed in Tracer and both the runs were combined in Log Combiner 1.8.3 with a 50% burn-in. Further, a maximum clade credibility tree was constructed keeping target heights in Tree Annotator 1.8.3 while allowing for a further 20% burn-in using a posterior probability limit of 0.95.

### 1.3 Ancestral area reconstruction

We have sought to understand the area of origin given their dispersal probabilities. We have reconstructed the ancestral area using a maximum likelihood method under the divergence– extinction–cladogenesis model (DEC) implemented in the software Lagrange build 20091004 (Ree & Smith, 2008). We have generated python scripts using the online configurator (http://www.reelab.net/lagrange) and run using python launcher.

Five areas were demarcated for examination of ancestral areas and the choice was based on the distribution of included taxa as described in Wilson (2011): A, Australia; B, Africa, and Madagascar; C, India; D, south-east Asia; E, South America (figure - 4). In addition, we have also retrieved the distribution data from other sources e.g. web-resource like Global Biodiversity Information Facility (GBIF), Encyclopaedia of Life (EOL), South African National Biodiversity Institute (SANBI), India Biodiversity Portal, to assign each genus to one or more areas (table-S5). We have restricted the number of ancestral areas were restricted to three given a wider cross-continental distribution of the family and to avoid added complexity of the analyses due to many areas. Our basic assumption was that the that past distributions of members of this group were similar to the present distribution. Given the medium dispersal ability of the extant members (mostly dispersal takes place through birds and small mammals like bats) of this group, we assumed that dispersal was possible only between adjacent areas (i.e. no dispersal was allowed between widely-separated areas e.g. A and B or E, and between C and E). Hence, the final permissible ancestral areas were ABC, ABD, AC, ACD, AD, BC, BCD, BCE, BD, BDE, BE, and CD. The relative position and connectivity among the five geographic areas underwent major change over the considered time period (110-0 Ma) i.e. post-Gondwana fragmentation and hence major events e.g. floatation of Indian landmass and its position relative to other landmasses, position of Afro-Madagascar and formation of transient land-bridges, and final collision into India were taken into account. Thus, we assigned dispersal probabilities between all adjacent areas within all time-slice based on geological history, climate history (Morley, 2000, 2003; Tiffney & Manchester, 2001; Zachos et al., 2001, Scotese, 2001), and general dispersal ecology of Myrtaceae, e.g. small to medium sized seeds means moderate level of dispersal, and grossly followed previous studies (Couvreur et al. 2011; Mao et al. 2012). After several exploratory runs with a range of dispersal probabilities and time-slices, we selected five dispersal probability category (low or no dispersal = 0.01; low dispersal = 0.25; medium dispersal = 0.5; high dispersal = 0.75; areas adjacent or very close = 1), and five time-slices (110-90, 90-65, 65-45, 45-30, and 30-0 Ma) for final model testing. Differences between models were assessed by directly comparing their respective log-likelihoods where the conventional cut-off value of two log-likelihood units considered to assess the statistical significance of likelihood differences (Ree et al. 2005). We evaluated the goodness of fit of the data for three alternative biogeographic models in an ML framework: model 0 was unconstrained with dispersal events between all adjacent areas possible during the whole period considered (110–0 Ma) which was considered as the null model. To account for Gondwana relict hypothesis model 1 was constrained in the following manner: all dispersal out of India were allowed with varied dispersal probability as predicted from past geologic events but restricting dispersal into India throughout 110Ma. Model 2: dispersal out of India was not allowed during 110 Ma as well as dispersal into India was not allowed till 45 Ma, but dispersal from south-east Asia during 45-0 Ma was allowed to emulate south-east Asian affinity (table-S6). We incorporated prior information on range evolution as well as probabilities between areas given discrete periods. Finally, we estimated the ancestral areas for all nodes relevant to this study.

## 2 Results

### 2.1 Phylogeny, molecular dating, and timing of divergence

A total of 45 sequences were generated for this study 15 each from three loci. Post-editing, the aligned dataset of 85 taxa consisted of 1824 sites with 923 variable, 583 parsimony informative, and 339 singleton sites. We scrutinised the resulting trees for weak nodal supports and collapsed such nodes (50% cutoff for bootstrapped nodes in RAxML and 0.9 cutoff for posterior probabilities in Bayesian reconstructions) in Tree Graph 2.11.1 (Stover and Muller 2010). All three methods of phylogenetic reconstructions revealed almost similar evolutionary relationships of major clades. In general, Myrtaceae *sensu lato* comprised two well-differentiated clades of subfamilies, Psiloxyloideae and Myrtoideae; subsequently, Myrtoideae split into tribes i.e. Lophostemoneae, Xanthostemoneae, Eucalypteae, Syncarpieae, Chamelaucieae, Leptospermeae, Syzygieae, Backhousieae, Metrosidereae, Tristanieae and Myrteae (figure-2a-b). However, the nodal supports for the lower division were low in the concatenated reconstructions of the partitioned data and could collapsed into a broad polytomy at which point topological incongruences between concatenation and species tree methods would disappear (figure-S1). Likewise, for shallower nodes, the partitioned concatenated reconstructions revealed better support and hence depicted monophyletic origin of the tribal clades *e.g.* Syzygieae, Myrteae, Eucalypteae, Leptospermeae, Chamelaucieae; of which Syzygieae and Myrteae were non-monophyletic in species tree. Whereas, in all three reconstructions, tribes Melaleuceae and Backhousieae portrayed monophyly. Within concatenated tree, Bayesian and ML tree revealed near similar topology. The minor differences were mostly due to comparatively lower nodal supports in the maximum likelihood tree compared to the Bayesian tree e.g. Metrosidereae tribe. Our tree demonstrated tribe-wise clades that was mostly in agreement with the previously published tress by Wilson et al. (2005), Biffin et al. (2010), and Thronhill et al. (2015) albeit some topological incongruences. Firstly, the position of the few tribes, e.g., Xanthostemoneae and Lophostemoneae that were sister to the rest of Myrtoideae, and Osbornieae and Melaleuceae that were sister to the rest of Myrtoideae except Xanthostemoneae, and Lophostemoneae, was other point of discordance. Secondly, other topological difference was in the position of Leptospermeae-Chamelaucieae clade that was sister to Eucalypteae in previous studies. These instances of discordance could be largely due to lack of power in out reconstructions since our reconstruction consisted of a subset of all taxa used in previous studies. However, our study focuses on the phylogenetic history of the Myrtaceae of WG of India and we are confident that these discordances would not affect our inferences.

**Figure 2:**
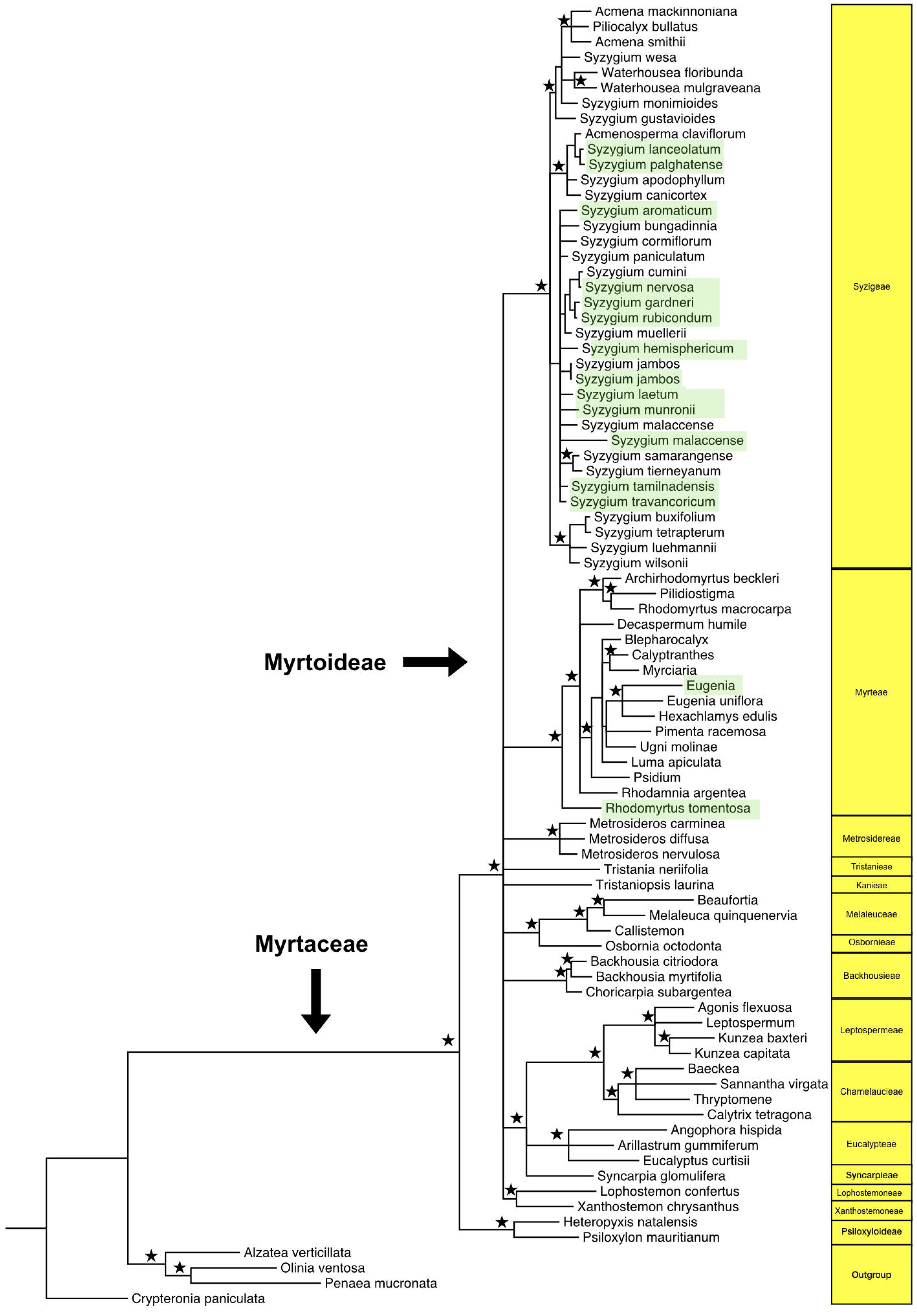
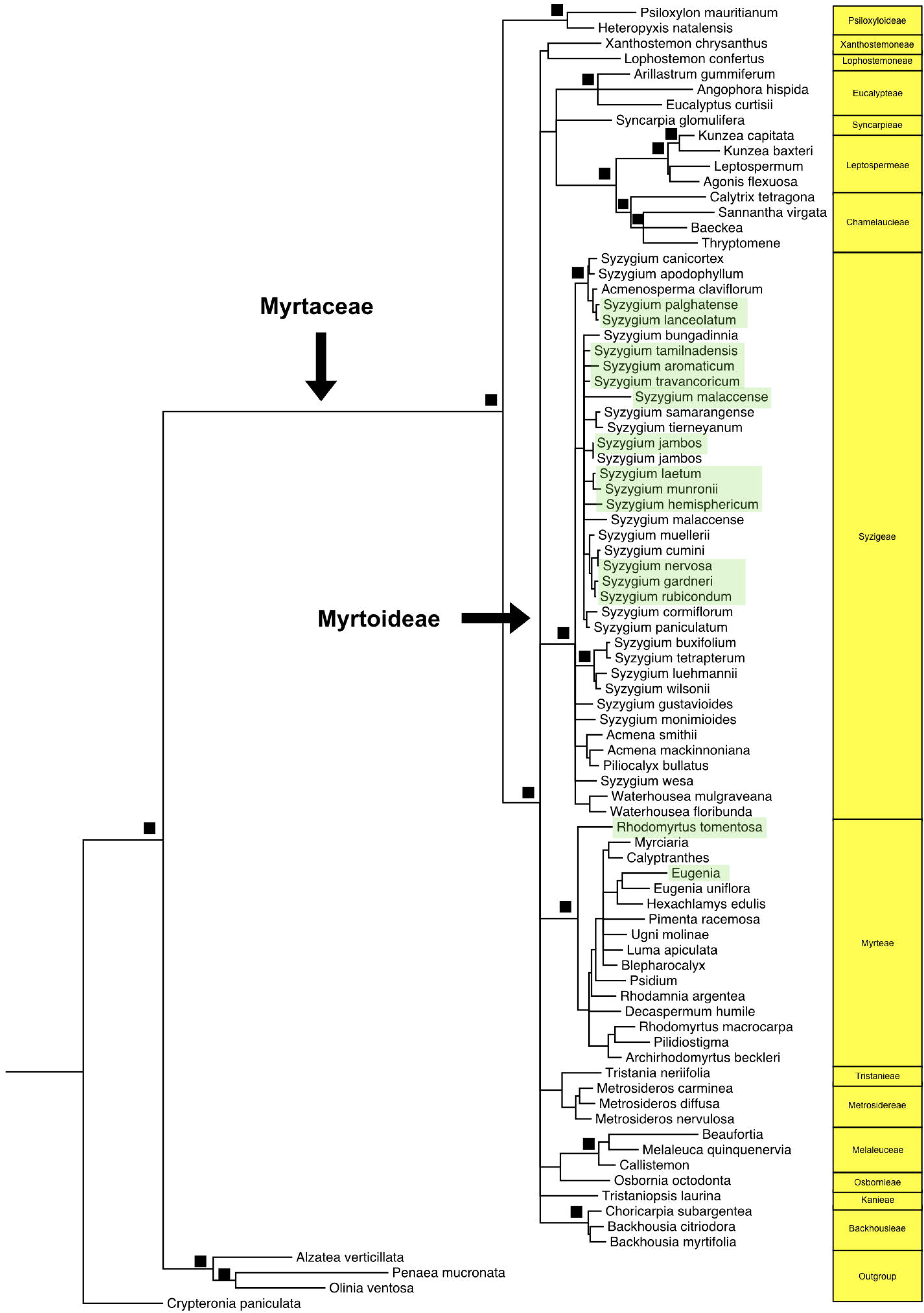
(a) Bayesian and (b) Maximum likelihood tree comprising 81-taxon of Myrtaceae and four outgroup taxa. Strongly supported nodes (posterior probability > 90%) are marked by stars in Bayesian tree. The closed squares at the nodes indicate high bootstrap support (>90%) in ML tree. Indian taxa are highlighted in light green.

The Fossil calibrated timetree portrayed that the oldest split between Myratceae *sensu lato* and outgroups occurred in the Cretaceous (Cenomanian-Turonian) c.a 95.7 Ma (95% HPD: 88.82 - 101.9 Ma) (figure-3, table - 2). Following this, two subfamilies of Myratceae *s.l.*i.e. Psiloxyloideae and Myrtoideae emerged in Santonian i.e. around 84.3 Ma (95% HPD: 77.93 - 90.15 Ma) with the subsequent origin of the most recent common ancestor (henceforth MRCA) of Myrtoideae happening in early Maastrichtian (68.83 Ma, 95% HPD: 63.11 - 73.07 Ma). Thereafter, clades of Myrtoideae continued rapid diversification in middle to late Paleocene (55.4 - 60.28 Ma). Most of the lower level cladogenesis were very recent, ranging from early to middle Miocene. Indian extant lineages appeared to be quite a young group born mostly during Miocene except *Rhodomyrtus* originating in Oligocene.

**Figure 3:**
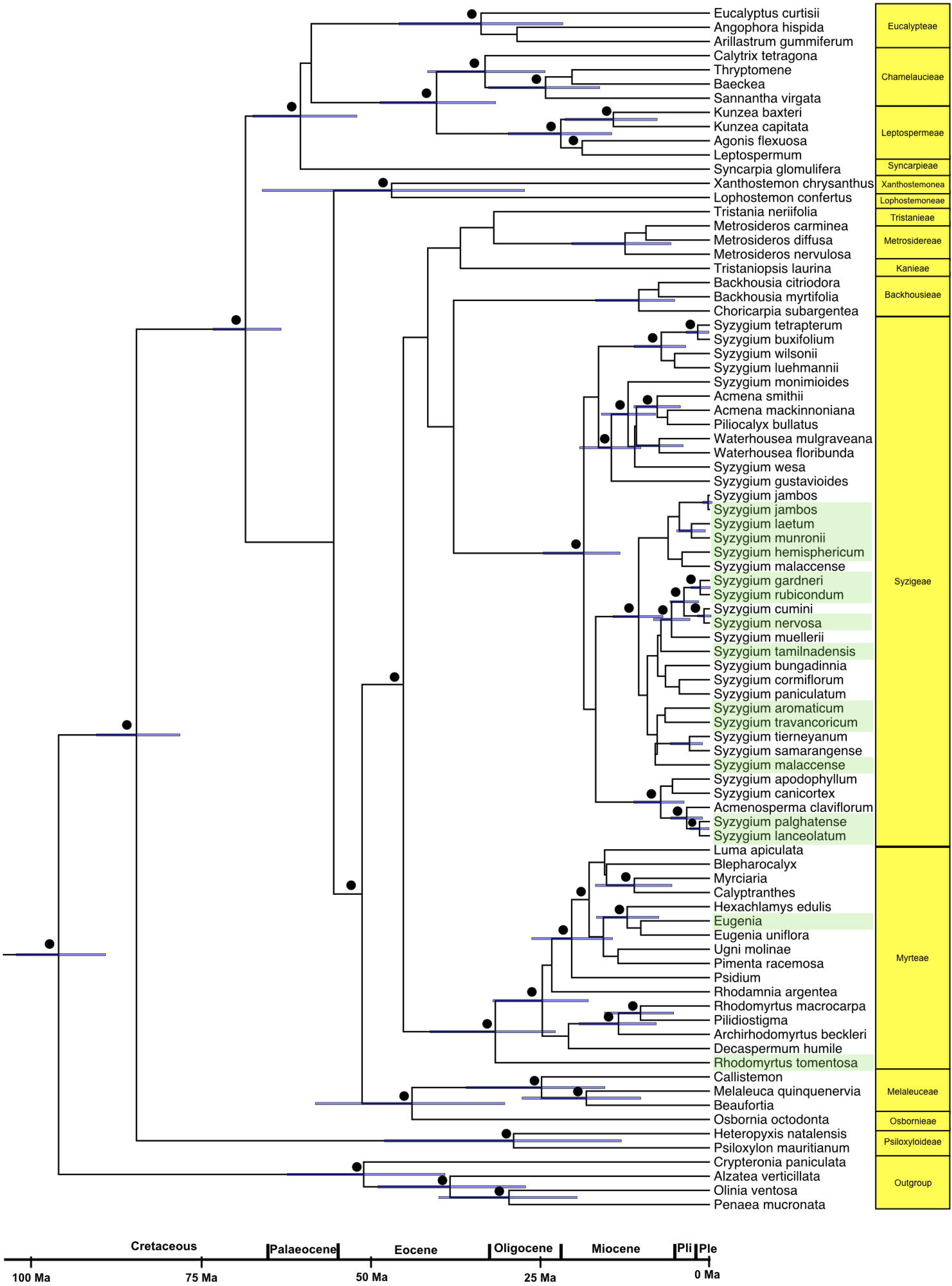
Bayesian maximum clade credibility tree of 85 taxa with four outgroup obtained from BEAST analyses. Nodes with blue node bars representing 95% highest posterior density intervals (Table 2). The numbers 1-4 represent four calibration points. Bayesian posterior probabilities < 0.9 are indicated by closed circles. A geological time scale is shown at the bottom with geological epoch abbreviations: Pli, Pliocene; Ple, Pleistocene.

**Figure 4:**
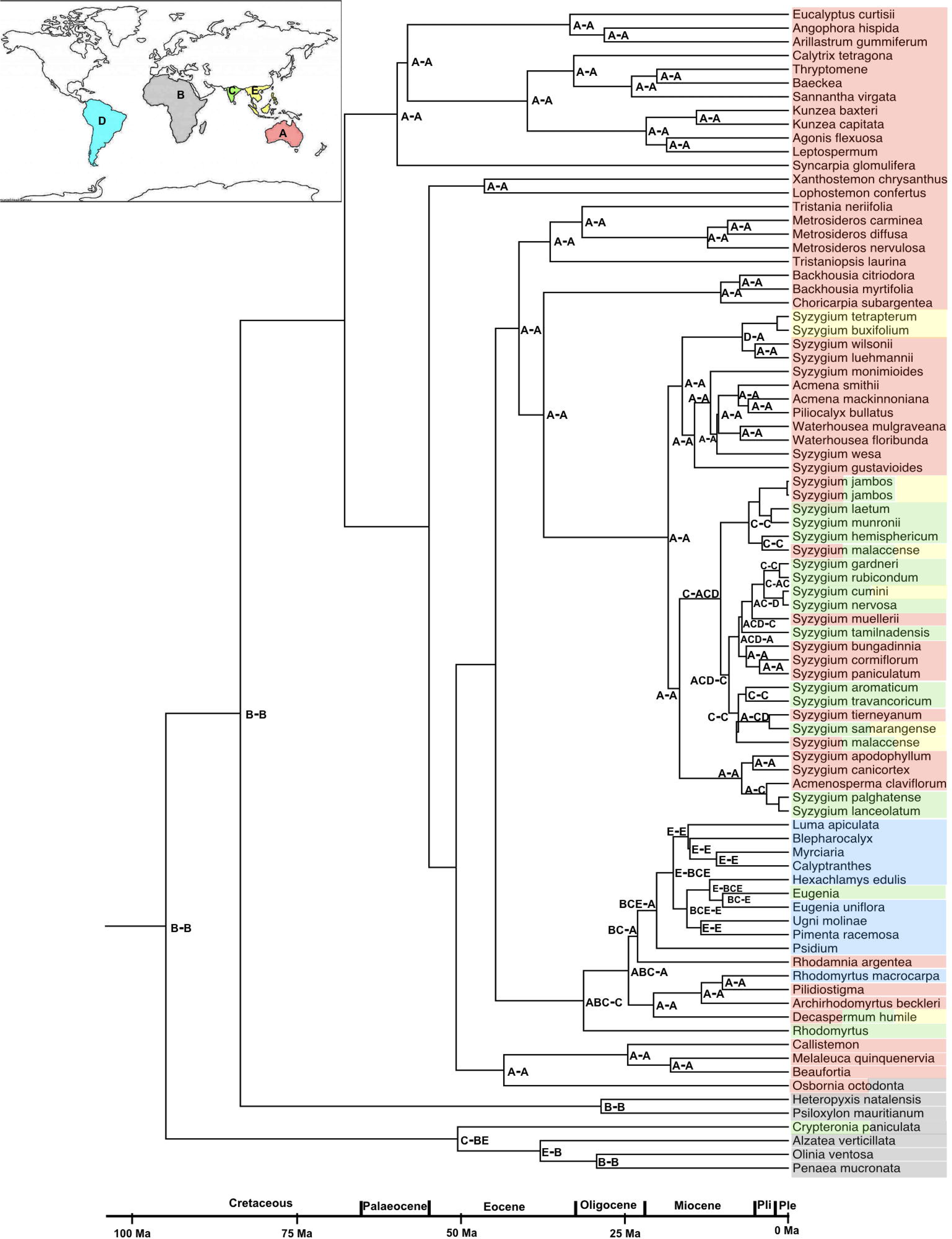
LAGRANGE derived cladogram of ancestral distributions in Myrtean lineages. Letters at nodes represent putative ancestral-area of the major nodes. The underscore separates the ancestral area reconstructed for the lower branch (left letter) from the one reconstructed for the upper branch (right letter) arising from the node. The letters at the node denote landmasses as following: A, Australia; B, Africa; C, India; D, Southeast Asia (without India); E, South America. Single-area letters indicate an ancestor restricted to a single area; two-letters indicate an ancestor restricted to two areas. The colours of the edges denote their current distribution and the same corresponds to the inset map: Light red - Australia, Light ash - Africa, Sky blue - South America, Light green - India, Yellow - south-east Asia. A geological time scale is shown at the bottom with geological epoch abbreviations: Pli, Pliocene; Ple, Pleistocene.

**Table-2:** Posterior age distributions (with 95% HPD) of major nodes of Myrtaceae

|  |  |
| --- | --- |
| Split between outgroup taxa and Myrtaceae<br>*sensu lato* | 95.7 Ma, 95% HPD: 88.82 - 101.9 Ma |
| MRCA of Myrtaceae | 84.3 Ma, 95% HPD: 77.93 - 90.15 Ma |
| MRCA of Myrtoideae | 68.33 Ma, 95% HPD: 63.12 - 73.07 Ma |

### 2.2 Ancestral area reconstruction

The reconstruction of biogeographic history revealed that the unconstrained model 0 (where dispersal to and from India was allowed throughout 110 Ma) outperformed other two models (table-3). Neither the Gondwana relict nor the south-east Asian dispersal model was a supported scenario. In summary, the best selected model suggested that the ancestors of outgroup taxa and Myrtaceae *s.l.* clade perhaps originated in the African landmass (a similar centre of origin was also depicted for the MRCA of Myrtaceae *s.l.* clade) (figure-4). Our analyses further indicated that the Austro-Africa-Indian landmass was the putative ancestral area for the Myrtoideae subfamily from where all the major genera have evolved. Afterwards, multiple dispersal events to Asia, South America, India and neighbouring island groups continued mostly during Eocene-Miocene. Origin of ancestors of *Syzygium* spp owed to Australia and dispersal to India was presumably mediated through the Asian mainland. Likewise, the Indian species of *Eugenia* portrayed a long distance dispersal from neotropics in Miocene. In contrast, *Rhodomyrtus* demonstrated origin in floating Indian landmass in Oligocene. However, it is noteworthy that many of the internal lower nodes lack strong posterior support thereby restricting us from making strong assumptions regarding their historical biogeography and future studies consisting of extensive taxon sampling as well as sequencing of more molecular markers may provide a better resolution in this regard.

**Table-3:** Results of ancestral area reconstruction using three different models of geographic range evolution (details of the models described in the text).

| Model | - loglikelihood<br>Ln(L) | Dispersal | Extinction |
| --- | --- | --- | --- |
| Model 0 (unconstrained dispersal) | 146.0 | 0.022 | 0.0007 |
| Model 1 (Gondwana element) | 152.6 | 0.021 | 0.0007 |
| Model 2 (south-east Asian affinity) | 152.7 | 0.022 | 0.0008 |

## 3 Discussion

Myrtaceae, a Gondwana family, is a key element of the south and south-east Asian tropical forests. The Western Ghats, a refuge of several taxa Myrtaceae, have been little explored to elucidate its history. Our central question on the origin of Myrtaceae diversity in the Western Ghats primarily sought to test two major hypotheses in historical biogeography, namely vicariance or dispersal. In the light of the interpretation of the evolutionary signatures of our molecular data, the extant flora seemed to be dispersed elements and not a derivative of Gondwana vicariance. However, our results do not entirely refute Gondwanan link as the vibrant fossil records in India are perhaps reminiscent of an extirpated Gondwanan flora.

### 3.1 Origin of the extant Myrtaceae lineages in WG

The shallow branches and recent dates of major clades of the sampled taxa indicated younger lineages of *Syzygium* and *Eugenia.* They presumably arrived via dispersal from south-east Asia or South America. On the other hand, *Rhodomyrtus tomentosa* appeared relatively older taxa, splitting early from rest of the Myrteae. It perhaps originated in floating Indian landmass and reached India via transoceanic dispersal.

The timing of MRCA of Crypteroniaceae, Peneaceae and Myrtaceae *s.l.*(95.7 Ma) roughly coincided with the completion of Gondwana break-up which echoed earlier inference (Thornhill et al. 2015). The dates of two oldest nodes (Myrtaceae stem and crown) in our study were very close to Biffin et al. (2010) (Myrtaceae stem (94 Ma) and crown (86 Ma)), and Thornhill et al. (2015) (Myrtaceae stem (91.6 Ma) and crown (85 Ma)) but different from Berger et al. (2016) who suggested relatively older dates for the Myrtaceae stem and crown while deciphering Myrtales biogeographic history. The different dates of Berger et al. (2016) possibly occurred due to dissimilar fossil calibration or incorporation of a large section of other taxa. The Eucalypteae stem node which was calibrated with an Indian fossil deviated widely from other studies (MRCA the tribe Eucalypteae showed 33.82 Ma in comparison to 40 Ma in Biffin et al. (2010) and 55.2 Ma in Thornhill et al. (2015).

Our sampled taxa mostly belonged to two tribes, i.e., Syzygieae and Myrteae. Myrteae, mostly a South American tribe, radiated during the Eocene to Miocene. According to our dated tree, the origin of Myrteae crown group happened in Oligocene (31.73 Ma) when Indian plate had collided with Asia and was staying far away from Africa-Australia-Antarctica. In contrast, early Myrteae diversification was shown to be 50.7 Ma (Thornhill et al. 2015), and 65.55 Ma (Vasconcelos et al. 2017 using macrofossil dates); however, the timing of Australian and South American Myrteae crown remained very close to our findings i.e. between 34.7 and 38.4 Ma (Thornhill et al. 2015). Near similar results were obtained by Biffin et al. 2010 (34 Ma) and by Vasconcelos et al (2017) (40.76 Ma) implementing microfossil dates. Within this clade, Indian species of *Rhodomyrtus* demonstrated an early split in Eocene-Oligocene transition perhaps while floating on Indian landmass. It is contradictory to south-east Asian origin of *Rhodomyrtus macrocarpa* which likely occurred in Oligocene-Miocene (Thornhill et al. 2015; Vasconcelos et al. 2017). It can be attributed to insufficient taxon sampling of *Rhodomyrtus* which is not a monophyletic genus with 18 species disjunctly distributed in southern China, eastern Australia and Caledonia (Snow et al. 2011, Wilson 2011). Majority of the species of *Eugenia* super-group is pantropical in distribution. Vasconcelos et al. (2017) have estimated the origin of the crown group of *Eugenia* in Oligocene with subsequent rapid radiation in Miocene. Most of such events had occurred in South and central America. Likewise, *Eugenia* portrayed an origin in late Miocene (10.42 Ma) in the clade populated with South American lineages probably suggesting a recent dispersal from neotropics. In Myrtaceae, many tribes including Myrteae produce succulent fruits which are devoured by birds and bats (Banack, 1998; Travest et al. 2001). Doing so, they may carry the seeds and promote long-distance dispersal and establishment of many taxa (henceforth LDDE).

The biogeographic evolution of *Syzygium,* a predominantly Australasian genus, appeared unequivocal from the past studies (Thornhill et al. 2015). LDDE has been cited as a likely cause of divergence between south-east Asian, Indian and Australian *Syzygium* (Thornhill et al. 2015). It is possible that multiple stepping stone colonisation from Australia via south-east Asia took place in the middle to late Miocene. A growing body studies support similar kind of dispersals from Australia to Asia (Barker et al., 2007; Renner et al., 2010). Those were facilitated by tectonic movements exerted by the collision of Australian plate with the Philippine and Asian plate. Dispersals from Australasia to Sunda over Oligocene-Miocene were likely mediated through i) island hop for well-dispersed taxa, ii) chance dispersals, and iii) dispersals of montane taxa (Morley 2003). A sudden increased abundance of Myrtaceae pollen around 17 Ma ascribed to the island hopping (Muller 1972). Many other plants, e.g., Podocarpaceae followed similar dispersal route to reach south-east Asia. Presumably, the dominance and the survival of rain-forest in Australia almost throughout its geologic history and its extension to the neighbouring island groups fuelled biotic migration (Morley, 2003). The dates of origin of Indian *Syzygium* fell appropriately in the time frame to support the view of overland dispersal from south-east Asia, probably via north-east Indian corridor. We can trace multiple such dispersals during the middle to late Miocene, but insufficient sampling and low resolvability of the markers have not deciphered the finer details.

Pertinent to say that Indian biological diversity owes significantly to the ‘into-India’ dispersal (Mani, 1974). Floral massing from south-east Asia by *Syzygium* seems to reiterate this emerging pattern. However, a general absence of empirical works impeded us to delineate its biogeographic significance and to mark it as a common dispersal route.

### 3.2 Alternate route: floatation on biotic ferry, colonization, and extinction

Although our molecular data supported relatively recent Neogene ‘into-India’ dispersal of extant lineages, it failed to reconcile with the abundance of Eocene to Miocene fossils in India. The fossil woods resembling *Eucalyptus* (*E. dharmendrae, E. ghughuensis, Eucalyptoxylon eocenicus*) of Maastrichtian-Danian in Deccan Intertrappean Bed strongly argue in favour of Myrtaceous presence on Indian Ferry (table - S1). Whereas, the oldest fossil of Australian *Eucalyptus* appeared in Eocene which is much younger compared Indian counterpart (Hermsen et al. 2010; Shukla et al. 2012). The prevalence Cenozoic fossils invoked an ancient origin tracing an alternate route after Gondwana break-up (figure - 1a-c).Perhaps, proto-Myrtaceae swam on the Indian plate, and/or arrived via early transoceanic exchange between landmasses via transient land bridges (Chatterjee & Scotese 1999; Briggs, 2003) e.g. in the mid-Early Cretaceous, when India was nearer to Australia-Antarctica (Birkenmajer and Zastawniak, 1986; Ali & Aichinson, 2008); and by Cenomanian, Turonian, and Early Coniacian, between African and Indian plate when Madagascar laid very close (Morley, 2003). Palynological records have provided strong support for such exchanges especially in plant families like Sapindaceae, Palmae, Myrtaceae (Srivastava 1987/88). This biotic flux gradually declined with India’s movement away from Madagascar towards mainland Asia (54-36 Ma). Verdant tropical rain forests during the drifting towards Asia created a conducive environment for the survival of several angiosperm families on Deccan plate e.g. persistence of Crypteroniaceae on Indian plate and subsequent out-of-India dispersal (Morley, 2000; Conti et al. 2002; Srivastava, 2011). It also draws further support from the rich dicot megafossils uncovered across a vast region of central and western India, which forms the Deccan Intertrappean Bed (Srivastava 2011). The flourishing tropical forest hinted at a climatic condition with heavy precipitation throughout the year quite similar to present day Western Ghats climate. The effect of tropical climate was also reflected in wood anatomical characters e.g. lack of growth rings, and ring porosity due to non-seasonality, simple perforated vessel elements, an absence of scalariform perforation plates owing to high rates of transpiration (Baas, 1982; Srivastava 2011).

However, the coupled effect of climate change in concert with Deccan volcanism had played a central role in Cretaceous-Paleogene mass extinction, including the disappearance tropical forest (Keller et al. 2009; Bajpai, 2009; Renne et al. 2015). The members of Myrtaceae along with several other angiosperm families presumably faced extirpation from Indian land except for the southern WG which served as refugia (Prasad et al. 2009). Similar extinction events had been quite common that tended to obscure robust interpretations. But, rich fossil records, despite an apparent absence of extant taxa, often invoked as the witness of existence in the past, e.g. fossils of marsupials, Rhynchocephalia, and plant families such as Araucariaceae and many Podocarpaceae in Northern Hemisphere (Crisp et al. 2011).

However, an absence of relevant information on various fronts e.g. the span and the intensity of climatic change, the spatial and temporal severity of the volcanism, and post-volcanism floral assemblage, constrained our inference. Several remaining questions can be elucidated with species-level well-resolved phylogeny. Thus, future research based on exhaustive sampling from India as well as from south-east Asia would allow delineating probable migration route(s), timing, and rate and potential driver(s) of the lineage diversification.

## Supporting information

a broad polytomy at which point topological incongruences between concatenation and species tree methods would disappear

elsewhere in India provides support to this Gondwana lineage hypothesis

## Data Availability

All the sequence generated in this study deposited in the Genbank (e.g. https://www.ncbi.nlm.nih.gov/nuccore/KT936455) and the corresponding accession numbers (KT936455, KT907476-77, KT936449-60, KT907475, KT907475, KT990096, KU049645-47, KU060784-86, KU060791-93, KU060787-90, KT970714-19, KT982668-76) are added in the supplementary information.

## Acknowledgements

We gratefully acknowledge the suggestions and assistance of the following persons: Vishnu Mukhri and Srikanth in field sampling; Kerala and Karnataka Forest Department for permitting collection of specimens. The project was primarily funded by SERB-fast track grant for young researchers (grant no. SERC/LS-158/2011). BC acknowledges fellowship and computational support from SEABIG grant (WBS R-154-000-648-646 and WBS R-154-000-648-733). Authors declare no competing interests.

## SUPPLEMENTARY INFORMATION

**Table S1**: A list of Myrtaceae fossils discovered in India

**Table S2**- Genbank accessions of the specimens used this study. * marked specimens are collected in this study.

**Table S3**- Voucher number of the specimens used this study

**Table S4**- ITS primer sequences used in this study

**Table S5:** Distribution matrix used in this study

**Table S6**: Dispersal matrix used to test various scenarios in LAGRANGE analyses. The letters represent as following: A, Australia; B, Africa; C, India; D, Southeast Asia (without India); E, South America. Probabilities of dispersal: 0.01, none or low; 0.25, medium-low; 0.5, medium; 0.75, medium-high; 1, high.

**Figure S1:** Bayesian species tree comprising 81-taxon of Myrtaceae and four outgroup taxa. Strongly supported nodes (posterior probability > 90%) are marked by triangles in Bayesian tree.

