## Supplementary figures and images for "Western Ghats Myrtaceae are not Gondwana elements but likely dispersed from south-east Asia"

### a broad polytomy at which point topological incongruences between concatenation and species tree methods would disappear

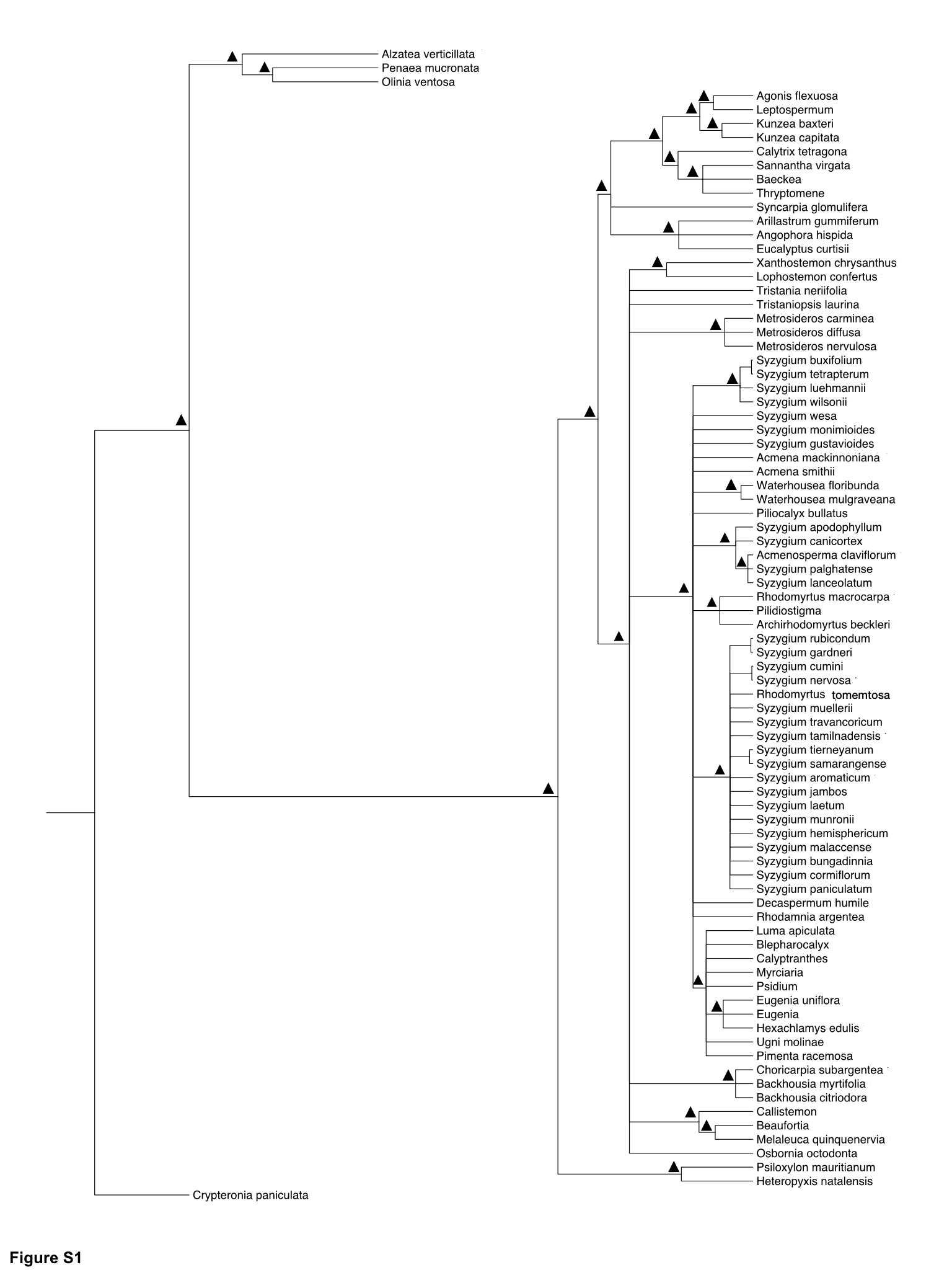
