## Supplementary material for "Western Ghats Myrtaceae are not Gondwana elements but likely dispersed from south-east Asia": elsewhere in India provides support to this Gondwana lineage hypothesis

Table S1: A list of Myrtaceae fossils discovered in India

| Specimen | Geological time period | Location | Reference |
| --- | --- | --- | --- |
| *Eucalyptus dharmendrae* | Maestrichtian-Danian (Early Tertiary) | Deccan Intertrappean Beds: Dindori (Mandla District) Madhya Pradesh | 1 |
| *Eucalyptus ghughuensis* | Late Maastrichtian–Danian | Deccan Intertrappean Beds: Ghughua near Shahpura on Shahpura–Niwas road, Dindori District, Madhya Pradesh | 2 |
| *Eucalyptoxylon vagadkholensis* | Palaeocene–early Eocene | Maljipura village, near Rajpardi lignite mine, Bharuch district, Gujarat | 3 |
| *Eucalyptoxylon eocenicus* | Eocene | Nal Clay Mine, Bikaner District, Rajasthan. | 3 |
| *Syzygioxylon chhindwarense* |  | Deccan Intertrappean Beds: near Mohagaon Kalan Locality in Chhindwara district, Madhya Pradesh | 4 |
| *Syzygium kasauliensis* | Lower Miocene | Kasauli Formation -Near Shiv Mandir on Kasauli-Jangeshu road, Solan District, Himachal Pradesh. | 5, 6 |
| *Syzygium nangwalbibrensis* | Upper Palaeocene | Tura Formation: Nangwalbibra near Williamnagar, East Garo Hills District, Meghalaya. | 6 |
| *Syzygium palaeobracteatum* | Middle Pliocene | Upper Siwalik: Bhikhnathoree, West Champaran District, Bihar | 7 |
| *Syzygium palaeocuminii* | Middle Miocene | Lower Siwalik: Sevok Road, near Teesta River bridge towards Oodlabari, Darjeeling District, West Bengal; | 8 |
|  | Miocene | Neyveli Lignite Deposit Mine I, South Arcot District, Tamil Nadu | 9 |
| *Callistemonoxylon decanensis* | Maastrichtian-Danian (Early Tertiary) | Deccan Intertrappean Bed | 10 |
| *Leptospermatoxylon indicum* |  |  |  |
| *Syzygium mandlaense* | Maestrichtian-Danian (Early Tertiary) |  |  |
| *Syzygium mohgaoense* |  |  |  |
| *Tristania confertoides* | Maestrichtian-Danian (Early Tertiary) |  |  |
| ?*Eugenia* sp. cf. *E americana* | Paleocene-Eocene | Gujarat |  |
| *Syzygium kachchhensis* | Paleocene-Eocene | Gujarat |  |

1 - **Bande MB, Mehrotra RC, Prakash U**. 1986 Occurrence of Australian element in the Deccan Intertrappean flora of India. *Palaeobotanist* 35: 1–12.

2 - **Shukla A, Mehrotra RC, Tyagi A.** 2012. The oldest fossil of Eucalyptus from the Late Maastrichtian–Danian of India and the theory of its Gondwanic origin. *Current Science* 103: 74-80

3 - **Shukla A, Mehrotra RC, Guleria JS.** 2014. Palaeophytogeography of *Eucalyptus* L'H'erit: New fossil evidences. *Journal of the Geological Society of India*, *84*(6), p.693-700

4 - **Nambudiri EMV, Tidwell WD.** 1977. *Syzygioxylon chhindwarense*, a new fossil wood from the Deccan Intertrappean series of India. *The Great Basin Naturalist*, pp.241-246.

5 - **Gulieria JS**. 2001. Fossil dicotyledonous woods from the Deccan Intertrappean beds of Kachchh, Gujarat, western India. *Palaeontographica Abteilung B*, pp.17-33.

6 - **Arya R, Guleria JS, Srivastava R.** 2001. New records of plant fossils from the Kasauli sediments of Himachal Pradesh, north-west India. *Phytomorphology*, *51*(1): 63-69.

7- **Awasthi N, Lakhanpal RN.** 1990. Additions to the Neogene florule from near Bhikhnathoree West Champaran District, Bihar. *Palaeobotanist* 37(3): 278-283.

8 - **Antal JS, Prasad M.** 1998. Angiospermous fossil leaves from the Siwalik sediments (Middle Miocene) of Darjeeling District, West Bengal. *Palaeobotanist* 46(3): 95-104.

9 - **Agarwal A.** 2002. Contributions to the fossil leaf assemblage from the Miocene Neyveli Lignite deposits, Tamil Nadu. *Palaeontographica* 261B: 167-206

10 - **Srivastava R.** 2011 Indian Upper Cretaceous-Tertiary Flora before Collision of Indian Plate: A Reappraisal of Central and Western Indian Flora. Memoir Geological Soc. India,77: 281-292

Table – S2: Genbank accessions of the specimens used this study, * marked specimens are collected in this study.

|  | Genbank accession number | | |
| --- | --- | --- | --- |
|  | *matk* | *ndhF* | *ITS* |
| *Acmena mackinnoniana* | DQ088543.1 | DQ088467 | AY187165 |
| *Acmena smithii* | DQ088545 | DQ088469 | AY187168 |
| *Acmenospermum claviflorum* | DQ088546 | DQ088470 | AY187169 |
| *Agonis flexuosa* | AF184711 | AY498762 | DQ499115 |
| *Alzatea verticillata* | AY151567 | AF215591 | AM235849 |
| *Angophora hispida* | AF368196 | AY498763 | AF190357 |
| *Archirhodomyrtus beckleri* | AF368197 | AY498764 | HQ225435 |
| *Arillastrum gummiferum* | AF368198 | AY498765 | AF058454 |
| *Backhousia citriodora* | AY525129 | AY498768 | KC134138 |
| *Backhousia myrtifolia* | AF368200.2 | DQ088472 | EF026609 |
| *Baeckea frutescens* | AF489365 | AY498770 | — |
| *Baeckea linifolia* | — | — | KM064974 |
| *Beaufortia sparsa* | KM065149 | — | KM064882 |
| *Beaufortia orbifolia* | — | AY498771 | — |
| *Blepharocalyx tweediei* | AY521531 | AY498772 | — |
| *Blepharocalyx salicifolius* | — | — | JN660936 |
| *Callistemon polandii* | AF184705 | AY498773 | — |
| *Callistemon viminalis* | — | — | JX856429 |
| *Calyptranthes pallens* | AF368201 | AY498775 | — |
| *Calyptranthes concinna* | — | — | AM234103 |
| *Calytrix tetragona* | AF489396 | AY498776 | KM064975 |
| *Choricarpia subargentea* | AF368202 | DQ088473 | EF026610 |
| *Crypteronia paniculata* | AB924733 | EU002217 | AM235848 |
| *Decaspermum humile* | AY521534 | AY498780 | AM234128 |
| *Eucalyptus curtisii* | AF368206 | AY498781 | AF390525 |
| *Eugenia sp** | KT907475 | KT990095 | KT970714 |
| *Eugenia uniflora* | AF368207 | DQ088457 | KM064994 |
| *Heteropyxis natalensis* | AF368208 | AY498824 | KM064805 |
| *Hexachlamys edulis* | AY525131 | AY498784 | KJ187652 |
| *Kunzea baxteri* | AF184722 | AY498789 | KM064868 |
| *Kunzea capitata* | AF184723 | AY498790 | EU833157 |
| *Leptospermum trinervium* | AF184735 | AY498792 | — |
| *Leptospermum laevigatum* | — | — | EU850664 |
| *Lophostemon confertus* | AF184707 | AY498794 | KM065037 |
| *Luma apiculata* | AY521540 | AY498795 | KM064889 |
| *Melaleuca quinquenervia* | GU135000 | EU410170 | AY835615 |
| *Metrosideros carminea* | AY521541 | AY498799 | KM064847 |
| *Metrosideros diffusa* | AY521542 | AY498800 | KM064992 |
| *Metrosideros nervulosa* | DQ088535 | DQ088458 | JF950781. |
| *Myrciaria vexator* | AY521544 | AY498804 | — |
| *Myrciaria floribunda* | — | — | AM234094 |
| *Olinia ventosa* | JX517344 | AF215594 | AM235855 |
| *Osbornia octodonta* | AF368213 | DQ088459 | EF041844 |
| *Penaea mucronata* | AY151589 | AF270756 | AM235871 |
| *Pilidiostigma sp* | AF368214 | AY498807 | — |
| *Pilidiostigma tropicum* | — | — | HQ225449 |
| *Pimenta racemosa* | AY521545 | AY498808 | AM234082 |
| *Psidium cattleyanum* | AB354959 | — | KM065002 |
| *Psidium guineense* | — | AY498810 | — |
| *Psiloxylon mauritianum* | AF368215 | AY498825 | EF026606 |
| *Rhodamnia argentea* | AF368217 | AY498810 | AM234129 |
| *Rhodomyrtus macrocarpa* | AY525137 | AY498811 | HQ225467 |
| *Rhodomyrtus tomentosa** | KT936455 | KT990096 | KT970715 |
| *Rhynchocalyx lawsoniides* | AF368218 | AF270757 | AM235850 |
| *Syygium apodophyllum* | DQ088558 | DQ088482 | AY187173 |
| *Syzygium aromaticum** | KT907476 | KU049645 | KT982668 |
| *Syzygium bungadinnia* | DQ088568 | DQ088490 | AY187182 |
| *Syzygium buxifolium* | DQ088569 | DQ088491 | KP093045 |
| *Syzygium canicortex* | DQ088570 | DQ088492 | AY187183 |
| *Syzygium cormiflorum* | DQ088572 | DQ088494 | AY187184 |
| *Syzygium cumini* | AY525140 | AY498814 | JF682812 |
| *Syzygium gardneri** | KT907477 | KU049646 | KT970719 |
| *Syzygium gustavioides* | DQ088582 | DQ088501 | AY187194 |
| *Syzygium hemisphericum** | KT936449 | KU049647 | KT982669 |
| *Syzygium jambos** | KT936450 | KU060786 | KT970716 |
| *Syzygium jambos* | DQ088583 | DQ088502 | KT970716 |
| *Syzygium laetum** | KT936451 | KU060785 | KT970717 |
| *Syzygium lanceolatum** | KT936452 | KU060784 | KT982670 |
| *Syzygium luehmannii* | DQ088587 | DQ088505 | AY187197 |
| *Syzygium malaccense** | KT936453 | KU060791 | KT982671 |
| *Syzygium malaccense* | DQ088590 | DQ088509 | AY187199 |
| *Syzygium monimioides* | DQ088544 | DQ088468 | AY187166 |
| *Syzygium muelleri* | DQ088593 | DQ088511 | EF026634 |
| *Syzygium munronii** | KT936454 | KU060792 | KT970718 |
| *Syzygium nervosa** | KT936456 | KU060793 | KT982673 |
| *Syzygium palghatense** | KT936457 | KU060787 | KT982674 |
| *Syzygium paniculatum* | DQ088598 | DQ088515 | AY187204 |
| *Syzygium rubicondum** | KT936458 | KU060788 | KT982675 |
| *Syzygium samarangense* | DQ088605 | AY498815 | KC815990 |
| *Syzygium tamilnadensis** | KT936459 | KU060789 | KT982672 |
| *Syzygium tetrapterum* | DQ088615 | DQ088527 | EF026649 |
| *Syzygium tierneynum* | DQ088616 | DQ088528 | AY187213 |
| *Syzygium travancoricum** | KT936460 | KU060790 | KT982676 |
| *Syzygium wesa* | DQ088617 | DQ088529 | AY187216 |
| *Syzygium wilsonii* | DQ088618 | DQ088530 | AY187219 |
| *Sannantha virgata* | EF581203 | EF581223 | KM064973 |
| *Syncarpia glomulifera* | AF368220 | AY498813 | KM065007 |
| *Thryptomene saxicola* | AF184709 | AY498816 | — |
| *Thryptomene calycina* | — | — | KM064883 |
| *Tristania nerifolia* | AF368224 | DQ088461 | EF026608 |
| *Tristaniopsis laurina* | AF184710 | AY498818 | KM064886 |
| *Ugni lominae* | AY525142 | AY498819 | KM064826 |
| *Waterhousea floribunda* | DQ088620 | DQ088531 | AY187221 |
| *Waterhousea malgraveana* | DQ088622 | DQ088533 | AY187223 |
| *Xanthostemon chrysanthus* | AF368227 | AY498823 | EF041515 |

**Table-S3**: Voucher number of the specimens used this study

| specimens | voucher number |
| --- | --- |
| *Eugenia sp** | HJCB-N-0303 |
| *Rhodomyrtus tomentosa** | HJCB-N-0302 |
| *Syzygium aromaticum** | HJCB- N- 0294 |
| *Syzygium gardneri** | HJCB-N- 110 |
| *Syzygium hemisphericum** | HJCB-N- 105 |
| *Syzygium jambos** | HJCB-N-0304 |
| *Syzygium laetum** | HJCB-N-0295 |
| *Syzygium lanceolatum** | HJCB- N- 0114 |
| *Syzygium malaccense** | HJCB-N-0298 |
| *Syzygium munronii** | HJCB-N- 103 |
| *Syzygium nervosa** | HJCB-N-0296 |
| *Syzygium palghatense** | HJCB-N-0297 |
| *Syzygium rubicondum** | HJCB-N-0301 |
| *Syzygium tamilnadensis** | HJCB-N- 104 |
| *Syzygium travancoricum** | HJCB-N- 107 |

Table S4: ITS primer sequence generated for this study:

| Code | sequence (5’———>3’) | Tm |
| --- | --- | --- |
| Sj-ITS-F | TCCTGCCTAGCAGAATGACC | 59.2 |
| Sj-ITS-R | GCTTAAACTCAGCGGGTAGC | 58.99 |

Table S5: Distribution matrix used in this study

|  | A | B | C | D | E |
| --- | --- | --- | --- | --- | --- |
| *Acmena mackinnoniana* | 1 | 0 | 0 | 0 | 0 |
| *Acmena smithii* | 1 | 0 | 0 | 0 | 0 |
| *Acmenosperma claviflorum* | 1 | 0 | 0 | 0 | 0 |
| *Agonis flexuosa* | 1 | 0 | 0 | 0 | 0 |
| *Alzatea verticillata* | 0 | 0 | 0 | 0 | 1 |
| *Angophora hispida* | 1 | 0 | 0 | 0 | 0 |
| *Archirhodomyrtus beckleri* | 1 | 0 | 0 | 0 | 0 |
| *Arillastrum gummiferum* | 1 | 0 | 0 | 0 | 0 |
| *Backhousia citriodora* | 1 | 0 | 0 | 0 | 0 |
| *Backhousia myrtifolia* | 1 | 0 | 0 | 0 | 0 |
| *Baeckea* | 1 | 0 | 0 | 1 | 0 |
| *Beaufortia* | 1 | 0 | 0 | 0 | 0 |
| *Blepharocalyx* | 0 | 0 | 0 | 0 | 1 |
| *Callistemon* | 1 | 0 | 0 | 0 | 0 |
| *Calyptranthes* | 0 | 0 | 0 | 0 | 1 |
| *Calytrix tetragona* | 1 | 0 | 0 | 0 | 0 |
| *Choricarpia subargentea* | 1 | 0 | 0 | 0 | 0 |
| *Crypteronia paniculata* | 0 | 0 | 1 | 1 | 0 |
| *Decaspermum humile* | 1 | 0 | 1 | 1 | 0 |
| *Eucalyptus curtisii* | 1 | 0 | 0 | 0 | 0 |
| *Eugenia* | 0 | 0 | 1 | 0 | 0 |
| *Eugenia uniflora* | 0 | 0 | 0 | 0 | 1 |
| *Heteropyxis natalensis* | 0 | 1 | 0 | 0 | 0 |
| *Hexachlamys edulis* | 0 | 0 | 0 | 0 | 1 |
| *Kunzea baxteri* | 1 | 0 | 0 | 0 | 0 |
| *Kunzea capitata* | 1 | 0 | 0 | 0 | 0 |
| *Leptospermum* | 1 | 0 | 0 | 0 | 0 |
| *Lophostemon confertus* | 1 | 0 | 0 | 0 | 0 |
| *Luma apiculata* | 0 | 0 | 0 | 0 | 1 |
| *Melaleuca quinquenervia* | 1 | 0 | 0 | 0 | 0 |
| *Metrosideros carminea* | 1 | 0 | 0 | 0 | 0 |
| *Metrosideros diffusa* | 1 | 0 | 0 | 0 | 0 |
| *Metrosideros nervulosa* | 1 | 0 | 0 | 0 | 0 |
| *Myrciaria* | 0 | 0 | 0 | 0 | 1 |
| *Olinia ventosa* | 0 | 1 | 0 | 0 | 0 |
| *Osbornia octodonta* | 1 | 0 | 0 | 1 | 0 |
| *Penaea mucronata* | 0 | 1 | 0 | 0 | 0 |
| *Pilidiostigma* | 1 | 0 | 0 | 0 | 0 |
| *Piliocalyx bullatus* | 1 | 0 | 0 | 0 | 0 |
| *Pimenta racemosa* | 0 | 0 | 0 | 0 | 1 |
| *Psidium* | 0 | 0 | 0 | 0 | 1 |
| *Psiloxylon mauritianum* | 0 | 1 | 0 | 0 | 0 |
| *Rhodamnia argentea* | 1 | 0 | 0 | 0 | 0 |
| *Rhodomyrtus sp* | 0 | 0 | 1 | 0 | 0 |
| *Rhodomyrtus macrocarpa* | 1 | 0 | 0 | 0 | 0 |
| *Sannantha virgata* | 1 | 0 | 0 | 0 | 0 |
| *Syncarpia glomulifera* | 1 | 0 | 0 | 0 | 0 |
| *Syzygium apodophyllum* | 1 | 0 | 0 | 0 | 0 |
| *Syzygium aromaticum* | 0 | 0 | 1 | 1 | 0 |
| *Syzygium bungadinnia* | 1 | 0 | 0 | 0 | 0 |
| *Syzygium buxifolium* | 0 | 0 | 0 | 1 | 0 |
| *Syzygium canicortex* | 1 | 0 | 0 | 0 | 0 |
| *Syzygium cormiflorum* | 1 | 0 | 0 | 0 | 0 |
| *Syzygium cumini* | 0 | 0 | 1 | 1 | 0 |
| *Syzygium gardneri* | 0 | 0 | 1 | 0 | 0 |
| *Syzygium gustavioides* | 1 | 0 | 0 | 0 | 0 |
| *Syzygium hemisphericum* | 0 | 0 | 1 | 0 | 0 |
| *Syzygium jambos* | 1 | 0 | 1 | 1 | 0 |
| *Syzygium jambos India* | 1 | 0 | 1 | 1 | 0 |
| *Syzygium laetum* | 0 | 0 | 1 | 0 | 0 |
| *Syzygium lanceolatum* | 0 | 0 | 1 | 0 | 0 |
| *Syzygium luehmannii* | 1 | 0 | 0 | 0 | 0 |
| *Syzygium malaccense* | 1 | 0 | 1 | 1 | 0 |
| *Syzygium malaccense India* | 1 | 0 | 1 | 1 | 0 |
| *Syzygium monimioides* | 1 | 0 | 0 | 0 | 0 |
| *Syzygium muellerii* | 0 | 0 | 0 | 1 | 0 |
| *Syzygium munronii* | 0 | 0 | 1 | 0 | 0 |
| *Syzygium nervosa* | 1 | 0 | 1 | 0 | 0 |
| *Syzygium palghatense* | 0 | 0 | 1 | 0 | 0 |
| *Syzygium paniculatum* | 1 | 0 | 0 | 0 | 0 |
| *Syzygium rubicondum* | 0 | 0 | 1 | 0 | 0 |
| *Syzygium samarangense* | 0 | 0 | 1 | 1 | 0 |
| *Syzygium tamilnadensis* | 0 | 0 | 1 | 0 | 0 |
| *Syzygium tetrapterum* | 0 | 0 | 0 | 1 | 0 |
| *Syzygium tierneyanum* | 1 | 0 | 0 | 0 | 0 |
| *Syzygium travancoricum* | 0 | 0 | 1 | 0 | 0 |
| *Syzygium wesa* | 1 | 0 | 0 | 0 | 0 |
| *Syzygium wilsonii* | 1 | 0 | 0 | 0 | 0 |
| *Thryptomene* | 1 | 0 | 0 | 0 | 0 |
| *Tristania neriifolia* | 1 | 0 | 0 | 0 | 0 |
| *Tristaniopsis laurina* | 1 | 0 | 0 | 0 | 0 |
| *Ugni molinae* | 0 | 0 | 0 | 0 | 1 |
| *Waterhousea floribunda* | 1 | 0 | 0 | 0 | 0 |
| *Waterhousea mulgraveana* | 1 | 0 | 0 | 0 | 0 |
| *Xanthostemon chrysanthus* | 1 | 0 | 0 | 0 | 0 |

Table - S6: Dispersal matrix used to test various scenarios in LAGRANGE analyses. The letters represent as following: A, Australia; B, Africa; C, India; D, south-east Asia (without India); E, South America. Probabilities of dispersal: 0.01, none or low; 0.25, medium-low; 0.5, medium; 0.75, medium-high; 1, high.

Model 0: Dispersal into and out of India allowed throughout the time period

0-30 Ma

|  | A | B | C | D | E |
| --- | --- | --- | --- | --- | --- |
| A | x | 0.01 | 0.01 | 0.75 | 0.01 |
| B |  | x | 0.25 | 0.25 | 0.01 |
| C |  |  | x | 1 | 0.01 |
| D |  |  |  | x | 0.01 |
| E |  |  |  |  | x |

30-45 Ma

|  | A | B | C | D | E |
| --- | --- | --- | --- | --- | --- |
| A | x | 0.01 | 0.01 | 0.75 | 0.01 |
| B |  | x | 0.25 | 0.25 | 0.01 |
| C |  |  | x | 1 | 0.01 |
| D |  |  |  | x | 0.01 |
| E |  |  |  |  | x |

45-65 Ma

|  | A | B | C | D | E |
| --- | --- | --- | --- | --- | --- |
| A | x | 0.01 | 0.01 | 0.25 | 0.01 |
| B |  | x | 0.01 | 0.25 | 0.25 |
| C |  |  | x | 0.75 | 0.01 |
| D |  |  |  | x | 0.01 |
| E |  |  |  |  | x |

65-90 Ma

|  | A | B | C | D | E |
| --- | --- | --- | --- | --- | --- |
| A | x | 0.01 | 0.25 | 0.01 | 0.01 |
| B |  | x | 0.25 | 0.25 | 0.01 |
| C |  |  | x | 0.25 | 0.01 |
| D |  |  |  | x | 0.01 |
| E |  |  |  |  | x |

90-110 Ma

|  | A | B | C | D | E |
| --- | --- | --- | --- | --- | --- |
| A | x | 0.5 | 0.75 | 0.01 | 0.75 |
| B |  | x | 1.0 | 0.01 | 1.0 |
| C |  |  | x | 0.01 | 0.75 |
| D |  |  |  | x | 0.01 |
| E |  |  |  |  | x |

Model 1: Dispersal into India not allowed throughout 110 Ma

0-30 Ma

|  | A | B | C | D | E |
| --- | --- | --- | --- | --- | --- |
| A | x | 0.01 | 0.01 | 0.75 | 0.01 |
| B | 0.01 | x | 0.01 | 0.25 | 0.25 |
| C | 0.01 | 0.25 | x | 1 | 0.01 |
| D | 0.75 | 0.25 | 0.01 | x | 0.01 |
| E | 0.01 | 0.25 | 0.01 | 0.01 | x |

30-45 Ma

|  | A | B | C | D | E |
| --- | --- | --- | --- | --- | --- |
| A | x | 0.01 | 0.01 | 0.75 | 0.01 |
| B | 0.01 | x | 0.01 | 0.25 | 0.25 |
| C | 0.01 | 0.5 | x | 0.75 | 0.01 |
| D | 0.75 | 0.25 | 0.01 | x | 0.01 |
| E | 0.01 | 0.25 | 0.01 | 0.01 | x |

45-65 Ma

|  | A | B | C | D | E |
| --- | --- | --- | --- | --- | --- |
| A | x | 0.01 | 0.01 | 0.01 | 0.01 |
| B | 0.01 | x | 0.01 | 0.25 | 0.01 |
| C | 0.25 | 0.5 | x | 0.25 | 0.01 |
| D | 0.01 | 0.25 | 0.01 | x | 0.01 |
| E | 0.01 | 0.01 | 0.01 | 0.01 | x |

90-65 Ma

|  | A | B | C | D | E |
| --- | --- | --- | --- | --- | --- |
| A | x | 0.5 | 0.01 | 0.01 | 0.75 |
| B | 0.5 | x | 0.01 | 0.01 | 1.0 |
| C | 0.5 | 0.5 | x | 0.01 | 0.75 |
| D | 0.01 | 0.01 | 0.01 | x | 0.01 |
| E | 0.75 | 1.0 | 0.01 | 0.01 | x |

110-90 Ma

|  | A | B | C | D | E |
| --- | --- | --- | --- | --- | --- |
| A | x | 0.75 | 0.01 | 0.01 | 0.75 |
| B | 0.75 | x | 0.01 | 0.01 | 1.0 |
| C | 0.75 | 1.0 | x | 0.01 | 0.75 |
| D | 0.01 | 0.01 | 0.01 | x | 0.01 |
| E | 0.75 | 1.0 | 0.01 | 0.01 | x |

Model 2: Dispersal out of India not allowed during 110 Ma, and dispersal into India not allowed during 110 - 45 Ma but allowed during 45 Ma

0-30 Ma

|  | A | B | C | D | E |
| --- | --- | --- | --- | --- | --- |
| A | x | 0.01 | 0.01 | 0.75 | 0.01 |
| B | 0.01 | x | 0.25 | 0.25 | 0.25 |
| C | 0.01 | 0.01 | x | 0.01 | 0.01 |
| D | 0.75 | 0.25 | 1 | x | 0.01 |
| E | 0.01 | 0.25 | 0.01 | 0.01 | x |

30-45 Ma

|  | A | B | C | D | E |
| --- | --- | --- | --- | --- | --- |
| A | x | 0.01 | 0.01 | 0.75 | 0.01 |
| B | 0.01 | x | 0.5 | 0.25 | 0.25 |
| C | 0.01 | 0.01 | x | 0.01 | 0.01 |
| D | 0.75 | 0.25 | 1 | x | 0.01 |
| E | 0.01 | 0.25 | 0.01 | 0.01 | x |

45-65 Ma

|  | A | B | C | D | E |
| --- | --- | --- | --- | --- | --- |
| A | x | 0.01 | 0.01 | 0.01 | 0.01 |
| B | 0.01 | x | 0.01 | 0.25 | 0.01 |
| C | 0.01 | 0.01 | x | 0.01 | 0.01 |
| D | 0.01 | 0.25 | 0.01 | x | 0.01 |
| E | 0.01 | 0.01 | 0.01 | 0.01 | x |

65-90 Ma

|  | A | B | C | D | E |
| --- | --- | --- | --- | --- | --- |
| A | x | 0.5 | 0.01 | 0.01 | 0.75 |
| B | 0.5 | x | 0.01 | 0.01 | 1 |
| C | 0.01 | 0.01 | x | 0.01 | 0.01 |
| D | 0.01 | 0.01 | 0.01 | x | 0.01 |
| E | 0.75 | 1 | 0.01 | 0.01 | x |

90-110 Ma

|  | A | B | C | D | E |
| --- | --- | --- | --- | --- | --- |
| A | x | 0.75 | 0.01 | 0.01 | 0.75 |
| B | 0.75 | x | 0.01 | 0.01 | 1.0 |
| C | 0.01 | 0.01 | x | 0.01 | 0.01 |
| D | 0.01 | 0.01 | 0.01 | x | 0.01 |
| E | 0.75 | 1 | 0.01 | 0.01 | x |
